# Distinct CA1 inputs support shifts in neural dimensionality and memory resolution

**DOI:** 10.64898/2026.01.14.699532

**Authors:** Cory McKenzie, Arna Ghosh, Adam I. Ramsaran, Bozhi Wu, Sakthivel Srinivasan, Tianwei Liu, Sheena A. Josselyn, Blake Richards, Paul W. Frankland

## Abstract

Event-based memories may be encoded with high (e.g., episodic) or low (e.g., gist) mnemonic resolution. While the CA1 region of the hippocampus encodes events at both scales, it is unclear how such dual coding emerges. Consistent with theoretical predictions^1,2^, here we show that trisynaptic (from CA3) and monosynaptic (from the medial entorhinal cortex) projections to CA1 encode high- and low-resolution event-based memories, respectively. Recruitment of the trisynaptic pathway during contextual fear conditioning promotes feedforward inhibition of CA1 via activation of parvalbumin-positive (PV^+^) interneurons. This allows sparse encoding of event-based memories in high-dimensional neural states that support high-resolution contextual fear memories, where mice exhibit conditioned fear only in the training context and not similar contexts. Inhibiting this input during training reduces activation of CA1 PV^+^ interneurons and feedforward inhibition, shifting neural dynamics to low-dimensional states that support low-resolution contextual fear memories where mice freeze in the training context and similar contexts. These experiments identify a circuit-based mechanism that causally links the dimensionality of CA1 neural dynamics to mnemonic resolution.

## Main text

While it is broadly accepted that the hippocampus encodes episodic memories (i.e., a high-resolution memory for an event tied to a specific context)^3–5^, there is emerging evidence that that the hippocampus may also encode gist memories (i.e., a low-resolution, event-based memory that lacks contextual details)^6–8^. At a behavioural level, episodic and gist memories offer complementary advantages for memory-based decision making^9,10^. High resolution, episodic memories permit perceptually-rich recollection, while minimizing interference with memories for closely-related situations (i.e., pattern separation^11^). Low resolution, gist memories avoid over-fitting to specific past situations, and can be used to guide behavior in more wide-ranging circumstances (i.e., generalization^12,13^). At a neural level, neural states that exist on high dimensional manifolds can represent more event features (or details) via orthogonalization, which helps to separate distinct representations. In contrast, non-orthogonal neural states that exist on low-dimensional manifolds are more likely to promote generalization^14–16^. Accordingly, episodic memories are hypothesized to depend on higher-dimensional neural representations that capture detailed features of the event. In contrast, gist memories are hypothesized to depend on compressed, low-dimensional neural representations, that capture coarser, potentially generalizable, features of an event^17^.

The hippocampus supports both high and low dimensional neural representation of task-relevant features^13,18,19^, and high dimensional neural dynamics in the human hippocampus correlate with episodic memory resolution^20,21^. How does the hippocampus achieve this dual coding ability? The CA1 region of the hippocampus receives indirect (via the trisynaptic pathway) and direct (via the monosynaptic pathway) input from the medial entorhinal cortex^22,23^. One influential theory proposes that these distinct inputs support high- and low-resolution encoding of event-based memories, respectively^1,2^. Here we tested this theory by selectively manipulating trisynaptic (CA3-to-CA1; Schaffer collateral) and monosynaptic (MEC-to-CA1; temporoammonic) projections to CA1 as mice formed an event (contextual fear) memory. We find that inhibition of the trisynaptic input during learning reduced mnemonic resolution, with mice exhibiting conditioned fear in both the training and similar contexts when subsequently tested. We show that trisynaptic projections preferentially contact CA1 PV^+^ interneurons, promoting feedforward inhibition. Combining calcium imaging with circuit manipulations in awake, behaving mice, we establish that loss of this feedforward inhibition, when trisynaptic projections are silenced, shifts neural states toward low dimensional neural representations resulting in low mnemonic resolution.

### CA3-to-CA1 projections required for high-resolution memory

The formation of episodic memory involves binding events to their surrounding spatial context^3^. This process may be modelled in mice using contextual fear conditioning where an aversive event (delivery of a footshock) becomes associated with a specific context^24^. In this task, mnemonic resolution may be assessed following conditioning by comparing freezing levels when mice are tested in the training apparatus (context A) versus a similar apparatus (context B), with freezing_A_>>freezing_B_ operationally defined as reflecting high mnemonic resolution as in previous studies^7,25–27^.

To explore how silencing distinct CA1 inputs impacts the resolution of contextual fear memories, we microinjected a retrogradely transported adeno-associated virus (AAV) expressing Cre recombinase into the dorsal CA1 (AAVrg-Cre), and AAVs expressing either a Cre-dependent inhibitory opsin (AAV-DIO-eNpHR3.0) or a control construct (AAV-DIO-eYFP) into CA3 or MEC (**Fig. 1a-d**). Implantation of optic fibres above CA3 and MEC allowed us to silence CA3-to-CA1 (i.e., trisynaptic) or MEC-to-CA1 (i.e., monosynaptic) projections to CA1, respectively, via delivery of red light photostimulation during contextual fear conditioning.

**Fig. 1.**
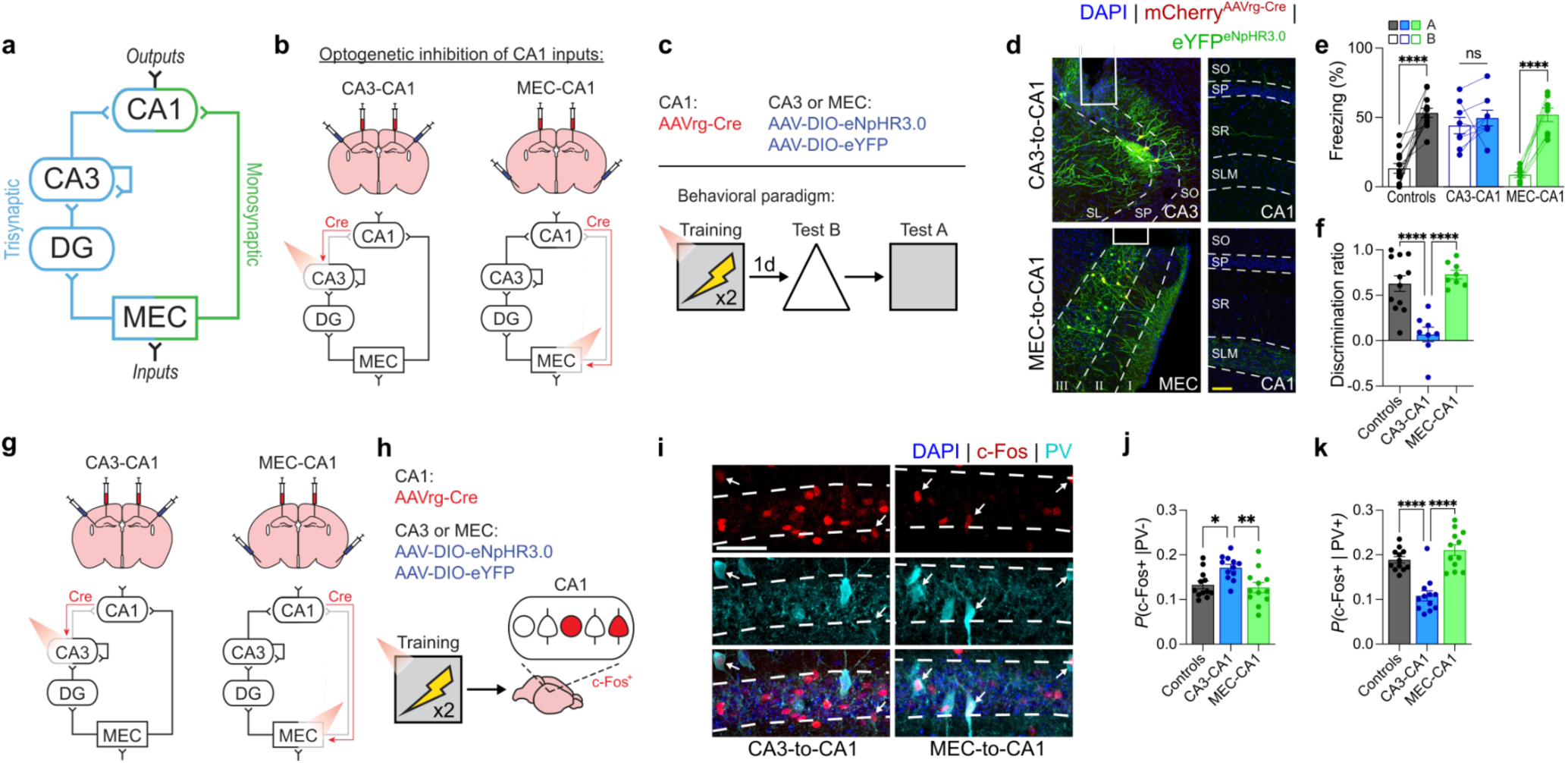
Silencing of CA3-to-CA1, but not MEC-to-CA1, projections reduces mnemonic resolution. **a)** Circuit diagram depicting dual inputs to CA1 originating in MEC via the trisynaptic (blue) or monosynaptic (green) projections. **b)** Experimental design for silencing trisynaptic and monosynaptic projections to CA1. AAVrg-Cre was injected into CA1 in all groups and AAV-DIO-eNpHR3.0 or AAV-DIO-eYFP was injected into either CA3 (trisynaptic expression of eNpHR3.0) or MEC (monosynaptic expression) to target trisynaptic and monosynaptic projections to CA1, respectively. **c)** Mice were contextually fear conditioned with red light inhibition during training. **d)** Left, representative images showing eNpHR3.0 expression in CA3 and MEC layer 3 with optic fibers implanted above. Right, labeled axons in CA1 with CA3 projections localizing to the SO and SR layers and direct MEC projections localizing to the SLM layer. **e)** Equivalent freezing in contexts A and B following CA3-to-CA1 projection silencing (two-way repeated measures ANOVA; Group x Context interaction: *F*_2,25_ = 13.16, *P* < 0.0001; n = 8 or 12 mice per group). **f)** Reduced mnemonic resolution following CA3-to-CA1 silencing reflected by decreased discrimination ratio, where discrimination ratio = [freezing_A_ – freezing_B_] / [freezing_A_+ freezing_B_] (one-way ANOVA; Group: *F*_2,25_ = 17.89, *P* < 0.0001; n = 8 or 12 mice per group). **g** and **h)** Experimental design to examine c-Fos induction following contextual fear training following CA3-to-CA1 or MEC-to-CA1 projection silencing. Viral strategy and training are the same as described in panel b, except mice are perfused 90 min after training and c-Fos in CA1 is quantified. **i)** Representative images of c-Fos and PV co-labelling in CA1 pyramidal layer (white arrows indicate co-labeled cells). **j)** CA3-to-CA1 inhibition increased c-Fos in PV^-^ cells (i.e., presumed pyramidal neurons) (one-way ANOVA; Group: *_F_*_2,33_ = 5.43, *P* = 0.0091; n = 12 mice per group). **k)** CA3-to-CA1 inhibition decreased c-Fos in PV^+^ interneurons (one-way ANOVA; Group: *F*_2,33_ = 26.59, *P* < 0.0001; n = 12 mice per group). Scale bars: white, 50 μm; yellow, 100 μm. Data are shown as mean ± SEM. \**P* < 0.05, \*\**P* < 0.01, \*\*\**P* < 0.001, \*\*\*\**P* < 0.0001. Lines link freezing scores across tests in the same mice.

Following training, control mice froze more in context A compared to context B (**Fig. 1e**), indicating that they formed high-resolution contextual fear memories. In contrast, when CA3-to-CA1 projections were optogenetically silenced during training, mice froze at equivalent levels in contexts A and B. Reduced discrimination between contexts A and B was not observed when MEC-to-CA1 projections were silenced (**Fig. 1f**), indicating that CA3-to-CA1, but not MEC-to-CA1, neurotransmission is important for the formation of high-resolution contextual fear memories. Similar loss of memory resolution occurred when trisynaptic (i.e., CA3-to-CA1), but not monosynaptic (i.e., MEC-to-CA1), projections to CA1 were chemogenetically silenced during training (**Extended Data Fig. 1**) or following region-specific optogenetic inhibition of any component exclusive to the trisynaptic pathway (**Extended Data Fig. 2**). Disabling both projections by optogenetically targeting MEC (i.e., a component common to both monosynaptic and trisynaptic pathways) reduced overall freezing levels in both contexts (**Extended Data 2**), indicating that at least one active input is necessary for memory formation.

Previous studies^25^ indicate that the formation of low-resolution contextual fear memories is associated with increased neuronal activation of CA1 during encoding. Therefore, we next tested whether silencing CA3-to-CA1 projections during contextual conditioning leads to an expansion of CA1 neural ensembles engaged by the learning experience. Using the same dual virus approach to optogenetically silence CA3-to-CA1 or MEC-to-CA1 projections during contextual fear conditioning (**Fig. 1g-h**), we evaluated training-induced CA1 activation by tracking induction of the activity-regulated gene, c-Fos (**Fig. 1i**). Silencing CA3-to-CA1, but not MEC-to-CA1, projections increased training-induced activation of CA1 (**Fig. 1j**).

Increased, rather than decreased, induction of c-Fos suggests that silencing excitatory CA3-to-CA1 projections leads to disinhibition of CA1 during training. Since CA1 activity is regulated by feedforward inhibition via PV^+^ interneurons^28,29^, we evaluated training-induced activation of PV^+^ interneurons with CA3-to-CA1 or MEC-to-CA1 projections silenced. Consistent with the hypothesis that silencing trisynaptic projections leads to a loss of feedforward inhibition^1^, training-induced c-Fos induction was reduced in PV^+^ interneurons when CA3-to-CA1, but not MEC-to-CA1, projections were silenced (**Fig. 1k**).

### CA3-to-CA1 projections preferentially contact PV^+^ interneurons

CA1 disinhibition following CA3-to-CA1 projection silencing raises the possibility that trisynaptic, rather than monosynaptic, projections preferentially innervate CA1 PV^+^ interneurons^30,31^ (**Fig. 2a**). To address this, we microinjected AAV1-Cre into either CA3 or MEC in Rosa-YFP reporter mice. Anterograde transneuronal spread of this virus results in Cre-induced recombination in post-synaptic target cells and YFP expression^32^ (**Fig. 2b**).

**Fig. 2.**
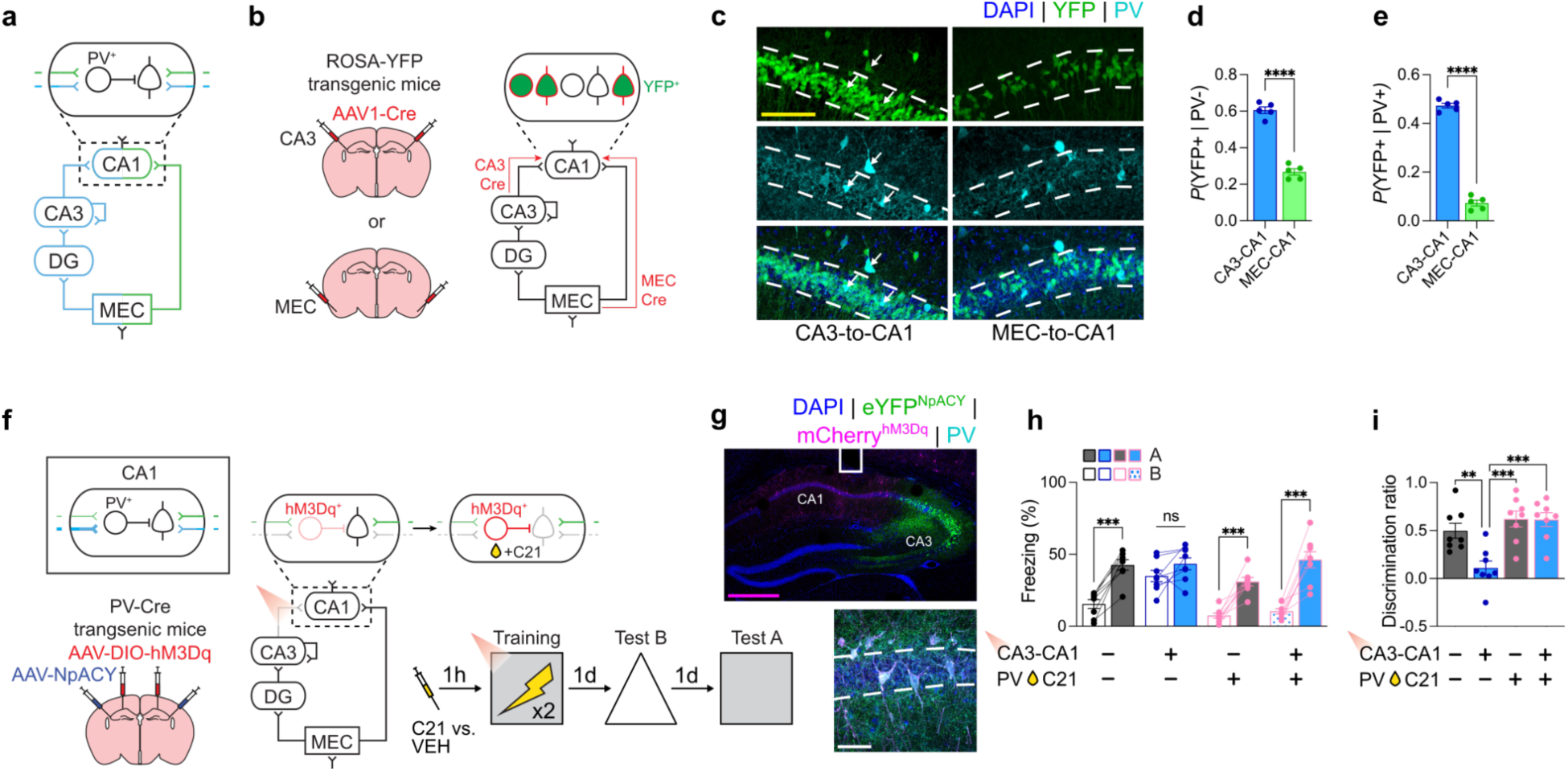
Mnemonic resolution depends on CA3 projections to PV^+^ interneurons in CA1. **a)** Circuit diagram highlighting cell-type specific inputs to CA1 from CA3 and MEC. **b)** Transsynaptic anterograde tracing was achieved by injecting AAV1-Cre in CA3 or MEC of ROSA-YFP transgenic mice. Cre protein expression in postsynaptic target neurons induces YFP expression. **c)** Representative images showing YFP and PV co-localization in CA1 (white arrows indicate co-labeled cells). **d)** PV^-^ CA1 neurons receive stronger input from CA3 compared to CA1 (unpaired *t* test; *t*_8_ = 13.46, *P* < 0.0001; n = 5 mice per group). **e)** CA1 PV^+^ interneurons receive 7-fold more input from CA3 than MEC (unpaired *t* test; *t*_8_ = 25.71, *P* < 0.0001; n = 5 mice per group). **f)** Experimental design to simultaneously activate PV^+^ interneurons in CA1 while inhibiting CA3-to-CA1 projections. AAV-DIO-hM3D_q_ was injected into CA1 of PV-Cre mice to induce expression of hM3D_q_ only in local PV^+^ interneurons. AAV-NpACY (containing an eNpHR3.0 opsin) was injected into CA3 and an optic fiber was placed over CA1 to inhibit CA3-to-CA1 projections with red light. **g)** Representative images showing NpACY expression in CA3 with axons in CA1, an optic fiber placed above CA1, and hM3D_q_ expression in PV^+^ interneurons in CA1. **h)** Activation of PV^+^ interneurons in CA1 restores mnemonic resolution following CA3-to-CA1 projection inhibition (three-way repeated measures ANOVA; Photostimulation x Drug x Context: *F*_1,28_ = 9.66, *P* = 0.0043; n = 8 mice per group). **i)** Discrimination ratios for experiment in f-h (two-way ANOVA; Photostimulation x Drug interaction: *F*_1,28_ = 6.26, *P* = 0.0185; n = 8 mice per group). Scale bars: white, 50 μm; yellow, 100 μm; magenta, 500 μm. Data are shown as mean ± SEM. \**P* < 0.05, \*\**P* < 0.01, \*\*\**P* < 0.001, \*\*\*\**P* < 0.0001. Lines link freezing scores across tests in the same mice.

Histological analyses identified YFP-tagged neurons in CA1 (**Fig. 2c**). Approximately twice as many CA1 PV^-^ neurons were YFP^+^ following CA3 compared to MEC microinjections, suggesting overall stronger input to CA1 from the trisynaptic compared to monosynaptic pathway (**Fig. 2d**). Notably, though, there were approximately 7-fold more PV^+^ interneurons that were YFP^+^ following CA3 compared to MEC microinjections (**Fig. 2e**). In contrast, a comparable number of SST interneurons were labeled following CA3 compared to MEC microinjections (**Extended Data Fig. 3a-c**). Red light inhibition of CA3-to-CA1 projections during contextual fear training reduced activation CA1 PV^+^, but not SST^+^, interneurons (**Extended Data Fig. 3d-g**), suggesting preferential innervation of CA1 PV^+^ interneurons by CA3-to-CA1 projections promotes feedforward inhibition and CA1 engram sparsification.

If silencing CA3-to-CA1 projections desparsifies CA1 encoding ensembles and reduces memory resolution via loss of PV^+^ interneuron-mediated feedforward inhibition, then artificially activating CA1 PV^+^ interneurons during training should restore mnemonic resolution. To test this, we combined optogenetic inhibition of CA3-to-CA1 projections with chemogenetic activation of CA1 PV^+^ interneurons during contextual fear conditioning. PV-Cre mice received microinjections of an AAV expressing a Cre-dependent G_q_-coupled Designer Receptors Exclusively Activated by Designer Drugs (DREADD), hM3D_q_ (AAV-DIO-hM3D_q_), into CA1, and an AVV expressing a red light sensitive inhibitory opsin (AAV-NpACY) into CA3 (**Fig. 2f-g**). Several weeks later mice were trained in the presence vs. absence of CA1 red light photostimulation (to inhibit CA3-to-CA1 terminals), and treated with the DREADD ligand, compound 21 (C21), or vehicle (VEH) (**Fig. 2f-g**).

Red light induced inhibition of CA3-to-CA1 projections during training reduced memory resolution, with mice freezing equivalently in contexts A and B when tested. Chemogenetic activation of CA1 PV^+^ interneurons had no effect alone, but was sufficient to restore memory resolution when combined with CA3-to-CA1 projection inhibition (**Fig. 2h-i**). These results support the idea that trisynaptic projections to CA1 promote memory resolution by increasing PV^+^ interneuron-mediated inhibition in CA1, resulting in sparser engram encoding.

### Inhibiting the trisynaptic pathway shifts CA1 to low dimensional coding

Our results indicate that parallel inputs to CA1 determine the resolution of contextual fear memories. Since orthogonalized neural states that sit on a high-dimensional manifold are capable of representing a larger set of distinct event features, we predicted that manipulation of these inputs during training would induce enduring shifts in the dimensionality of neural representations corresponding to the learned experience. Mice expressing a genetically-encoded calcium indicator (Thy1-GCaMP6f mice) were implanted with miniature fluorescence endoscopes to image CA1 neuronal activity^33^. To inhibit trisynaptic or monosynaptic projections to CA1, we microinjected a retrogradely transported AAV expressing Cre recombinase into the dorsal CA1 (AAVrg-Cre), and AAVs expressing either a Cre-dependent inhibitory DREADD (AAV-DIO-hM4D_i_) or a control construct (AAV-DIO-eYFP) into CA3 or MEC (**Fig. 3a-b**).

**Fig. 3.**
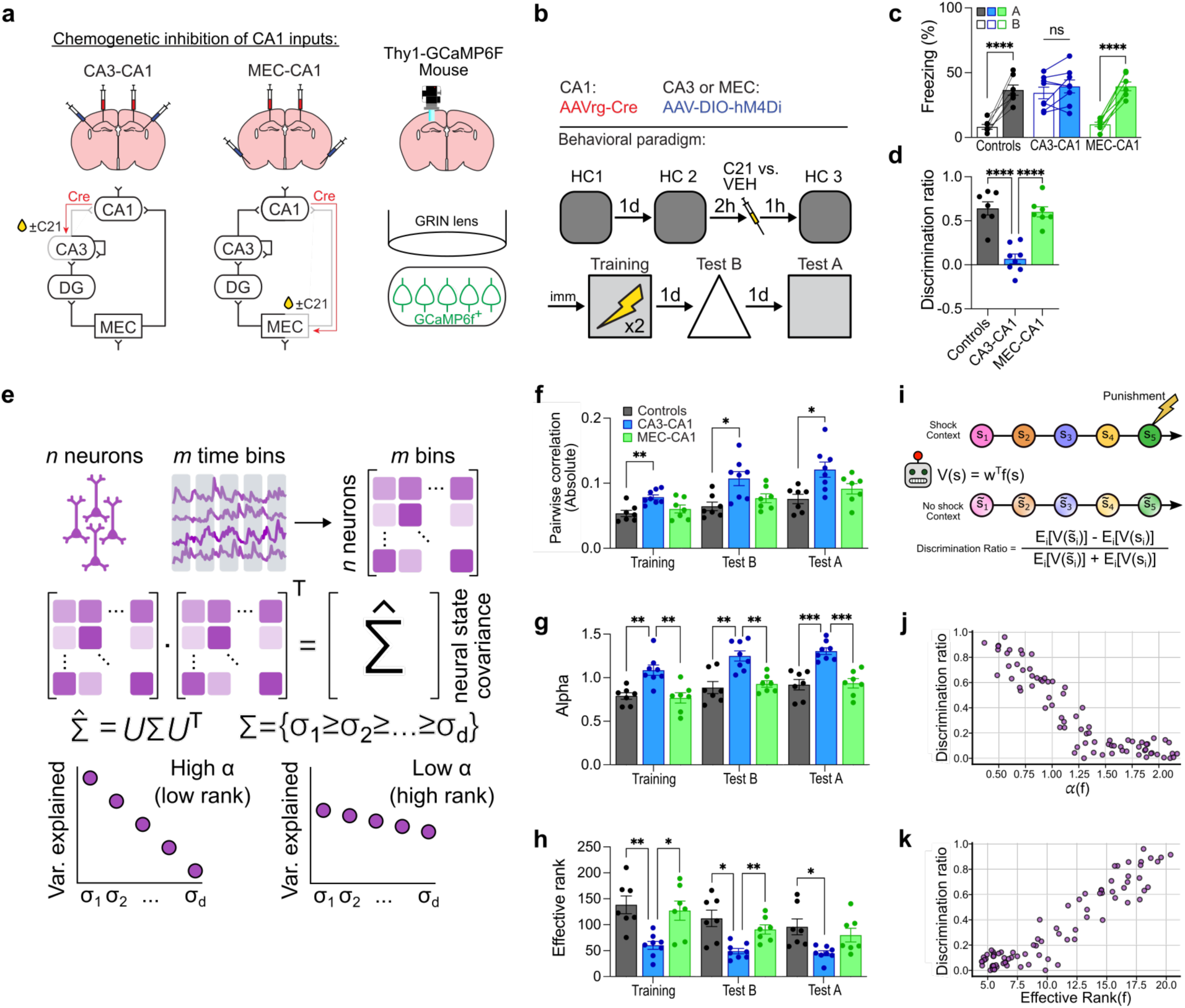
Inhibition of CA3-to-CA1 projections reduces the dimensionality of CA1 memory representations. **a)** Experimental design for chemogenetically inhibiting CA3-to-CA1 or MEC-to-CA1 projections in combination with 1-photon calcium imaging in CA1. In Thy1-GCaMP6F mice AAVrg-Cre was injected into CA1 and AAV-DIO-hM4D_i_ was injected into either CA3 or MEC. A GRIN lens was implanted unilaterally above CA1, and mice were given 4-6 weeks to recover. **b)** CA1 calcium transients were imaged in 3 homecage (HC) sessions (5 min each). C21 or VEH control was injected an hour before HC3 and training (5 min preshock, 2 footshocks 1min apart, and 5min postshock), and on subsequent days while mice were tested in contexts A and B (10 mins each). **c)** CA3-to-CA1 projection inhibition resulted in equivalent freezing in contexts A and B (two-way repeated measures ANOVA; Group x Context interaction: *F*_2,19_ = 14.83, *P* < 0.0001; n = 7 or 8 mice per group). **d)** Reduced discrimination following CA3-to-CA1 projection inhibition (one-way ANOVA; Group: *F*_2,19_ = 27.37, *P* < 0.0001; n = 7 or 8 mice per group). **e)** Neural activity cross-covariance matrices were used to assess alpha and rank measures of dimensionality. High alpha values (faster power law decay of the eigenspectrum) and low effective rank are associated with low dimensionality. **f)** CA3-to-CA1 inhibition increased pairwise correlations between all neuron pairs (two-way repeated measures ANOVA; Group: *F*_2,19_ = 9.93, *P* = 0.0011; n = 7 or 8 mice per group). **g)** Inhibiting CA3-to-CA1 projections increased alpha values in all sessions (two-way repeated measures ANOVA; Group: *F*_2,19_ = 22.57, *P* < 0.0001; n = 7 or 8 mice per group). **h)** Inhibiting CA3-to-CA1 projections reduced rank across all sessions (two-way repeated measures ANOVA; Group: *F*_2,19_ = 9.56, *P* = 0.0013; n = 7 or 8 mice per group). **i)** A computational model was set up to empirically assess how dimensionality affects the discrimination between a shock and a neutral context. The context representations were optimized with gradient descent to achieve a specific dimensionality and the agent attempts to discriminate between the contexts. **j)** Modelling predicted that higher alpha values permitted less efficient discrimination between contexts (spearman correlation; ρ = - 0.87, P < 0.0001). **k)** Higher rank permitted more efficient discrimination between contexts (spearman correlation; ρ = 0.86, P < 0.0001). Data are shown as mean ± SEM. \**P* < 0.05, \*\**P* < 0.01, \*\*\**P* < 0.001, \*\*\*\**P* < 0.0001. Lines link freezing scores across tests in the same mice.

Chemogenetic inhibition of CA3-to-CA1, but not MEC-to-CA1, projections during training resulted in low resolution, contextual fear conditioning memories with mice freezing equivalently in contexts A and B (**Fig. 3c-d**), replicating our previous experiments (**Fig. 1e-f**, **Extended Data Figs. 1-2**). CA1 population activity was tracked in home cage sessions prior to training (**Fig. 3e**), and during subsequent training and testing (4649 neurons from 7 control mice, 4134 neurons in 8 mice in the CA3-to-CA1 inhibition group, and 4281 neurons in 7 mice in the MEC-to-CA1 inhibition group). After removing spontaneous home cage activity to isolate task-relevant activity (see Methods and **Extended Fig. 5**), we observed elevated CA1 activation following CA3-to-CA1, but not MEC-to-CA1, inhibition, consistent with our c-Fos analyses (**Extended Data 5**).

To explore the statistical structure of the neural states in CA1 that emerge through conditioning we computed pairwise correlations for all neuron pairs. Silencing CA3-to-CA1, but not MEC-to-CA1, projections increased the magnitude of pairwise correlations relative to controls during training, and during subsequent testing in contexts A and B (**Fig. 3f**). This raises the possibility that manipulating trisynaptic projections alters the dimensionality of the manifolds on which CA1 task-relevant neural states lie. Therefore, we computed the decay rate of the power spectrum of the bootstrapped neural activity cross-covariance matrix (i.e., alpha), which provides a measure of the dimensionality and smoothness of neural manifolds^15^ (see Methods) (**Fig. 3e**). Inhibiting CA3-to-CA1, but not MEC-to-CA1, projections during training increased the alpha value of CA1 neural representations indicating a decrease in dimensionality and an increase in smoothness of the manifolds, in-line with decreased orthogonality (**Fig. 3g**). To further verify this change in neural representational geometry we measured the effective rank of the cross-correlation matrix, which provides an alternate measure of dimensionality^34^. In agreement with the changes in alpha, inhibiting CA3-to-CA1 projections, but not MEC-to-CA1 projections, resulted in a reduction of the effective rank of the representational manifolds during training and testing (**Fig. 3h**). These shifts in dimensionality were observed during training (i.e., during chemogenetic inhibition of CA3-to-CA1 projections) and also during subsequent testing in contexts A and B (i.e., in absence of the DREADD ligand, C21), indicating that inhibition of trisynaptic projections during learning induces enduring task-relevant changes in representational geometry. Moreover, dimensionality shifts were specific to learning since similar chemogenetic inhibition of CA3-to-CA1 projections did not alter the dimensionality of CA1 neural representations at baseline (i.e., in home cage) (**Extended Data Fig. 7**).

To better understand how changes in the dimensionality of the manifolds might affect behavior, we built a computational model of an agent learning to associate two different contexts with different valence. We manipulated the dimensionality of the context representations using gradient descent (see Methods) and calculated the ability of the agent to discriminate between a shock context and a neutral context (**Fig. 3i**). We observed that representations with a higher alpha and lower effective rank, as observed following CA3-to-CA1 inhibition, supported less efficient discrimination between contexts (**Fig. 3j-k**). Thus, the changes we observed in the dimensionality of CA1 neural representations following inhibition of the trisynaptic pathway should, according to computational principles, reduce discrimination, as we observed behaviourally (**Fig. 3c-d**).

## Discussion

The hippocampus can support high- or low-resolution event-based memories^6,7,25,35^, but it is unclear how this computational trade-off is accomplished. Our experiments identify a circuit-based mechanism that determines mnemonic resolution by regulating the dimensionality of CA1 neural dynamics. Engagement of trisynaptic projections to CA1 during training promotes feedforward inhibition via activation of CA1 PV^+^ interneurons, promoting orthogonalized, high-dimensional CA1 neural dynamics that support high-resolution memories of an experienced event. In contrast, in the absence of trisynaptic projections, engagement of monosynaptic projections to CA1 during training results in weaker PV^+^ interneuron-mediated feedforward inhibition, promoting non-orthogonal, low-dimensional CA1 neural dynamics that support low-resolution memories of an experienced event.

Consistent with the Complementary Learning Systems (CLS) model^36^, we found that the trisynaptic pathway is the default circuit mechanism for event memory formation, with engagement of the trisynaptic pathway supporting the rapid formation of high-resolution memory representations in CA1. While the CLS model emphasizes that excitatory connectivity between regions of the hippocampus supports the computations (e.g., pattern separation) necessary for forming high-resolution, episodic memories, our results identify CA3-to-CA1 PV^+^ interneuron-mediated feedforward inhibition as the key factor for orthogonalizing memory representations for events. Although CA1 is strongly influenced by CA3 projections to both its principal cells and PV^+^ interneurons^31^, the net effect of these projections is to constrain the size of CA1 neuronal ensembles during learning.

The CLS model proposes that generalized memory representations are slowly acquired by the cortex. In contrast, our experiments indicate that monosynaptic MEC-to-CA1 projections are sufficient for this type of learning. Our findings are therefore consistent with more recent computational models of hippocampal memory formation^1,2^ in which parallel trisynaptic and monosynaptic hippocampal pathways encode specific (episodic) and generalized (gist) memories, respectively. Our experiments further reveal that generalized memory representations are formed by MEC-to-CA1 projections because, unlike CA3-to-CA1 projections, these only weakly engage PV-mediated feedforward inhibition mechanisms during learning. That the hippocampus encodes specific memories under normal conditions suggests that the trisynaptic pathway is the predominant circuit for forming event memories, analogous to the predominance of the hippocampus, rather than cortex, theorized by CLS^36^.

The influence of the monosynaptic pathway on memory resolution and CA1 dimensionality may be more pronounced at earlier developmental stages and/or in disease states. Importantly, reduced PV^+^ interneuron-mediated inhibition is a shared feature of these states. For example, PV^+^ interneurons are functionally immature in CA1 before the fourth postnatal week, and do not support feedforward inhibition during memory encoding^25^. Accumulation of extracellular matrix structures known as perineuronal nets in CA1 during the fourth postnatal week stabilizes excitatory projections onto PV^+^ interneurons^37,38^ enabling adult-like feedforward inhibition and the emergence of high-dimensional memory representations. Accordingly, breakdown of PNNs and the consequent retraction of excitatory drive onto hippocampal PV^+^ interneurons in normal and pathological aging (e.g., Alzheimer’s disease) impairs the function of the trisynaptic pathway to encode detailed memories^39–43^.

The ability to simultaneously track activity of many neurons has made it possible to visualize the representational geometry of population activity in different brain regions, in different species and during different behaviors^15,16,20,44–46^. Within the hippocampus, for example, increasing dimensionality of neural representations is correlated with increasing mnemonic resolution^13,18^. Conversely, low dimensional neural states correspond to more abstracted representations of task-relevant features, allowing for inference and generalization^21^. While these studies indicate that the hippocampus can support both low and high dimensional neural representation of task-relevant features, here we identified circuit-based manipulations that are sufficient to cause shifts in the dimensionality of CA1 neural states and that these lead to corresponding shifts in mnemonic resolution. These results are in line with previous predictions that CA1 supports both high and low mnemonic resolution by integrating information from both CA3 and the entorhinal cortex^2^ in order to balance mnemonic precision and flexibility^13^.

## Methods

### Mice

Four mouse lines were used. Male and female F1 hybrid (C57BL/6NTac x 129S6/SvEvTac) wild-type (WT) mice were used for optogenetic, chemogenetic, and c-Fos based experiments unless otherwise stated. Rosa-YFP mice (B6;pNestinCRExROSA26YFP) which express the YFP protein in the presence of Cre recombinase were used for anterograde tracing^47^. Rosa-YFP breeders were maintained on a C57BL/6 background and crossed with 129S6/SvEvTac breeders to generate the hybrid Rosa-YFP mice used in experiments. PV-Cre knock-in driver transgenic mice (B6;129P2-Pvalbtm1(cre)Arbr/J, Strain# 008069), which express Cre recombinase in PV^+^ interneurons without disrupting endogenous PV expression, were originally generated by Silvia Arber (FMI)^48^ and were obtained from Jackson Laboratory. These mice were used to test the effect of combined PV^+^ interneuron activation with CA3-to-CA1 projection inhibition. Homozygous PV-Cre mouse breeders were maintained on a C57BL/6 background and crossed with 129S6/SvEvTac breeders to generate the hybrid PV-Cre mice used in experiments. Thy1-GCaMP6f mice were used in the calcium imaging experiment. We obtained mice hemizygous for the Thy1-GCaMP6f transgene^49^ on a C57BL/6 background (GP5.17 line, Jackson Laboratories) and crossed these with 129S6/SvEvTac mice. The resulting offspring were on a hybrid background, similar to the WT mice used in previous experiments. Only offspring hemizygous for the GCaMP6f transgene were used for the GCaMP imaging experiment. Mice were bred at the Hospital for Sick Children and group-housed on a 12-h light/dark cycle with food and water ad libitum. All experiments took place during the light phase. All procedures were approved by the Hospital for Sick Children Animal Care and Use Committee and conducted in accordance with Canadian Council on Animal Care and National Institutes of Health guidelines.

### Drugs

*DREADD agonist 21 (C21).* C21 dihydrochloride (Tocris, cat# 6422) was prepared as a stock solution of 10 mg/ml in dH_2_O and stored at -20 °C. Stock solution was later thawed and diluted 1:10 in PBS. Diluted C21 was administered via i.p. injection 1 h before contextual fear training (1.0 mg/kg) to activate or inhibit DREADD-expressing neurons.

### Adeno-associated viruses (AAVs)

All AAVs, apart from AAV1-Cre, were made in house.

We used adeno-associated viruses to manipulate neuronal activity in a circuit specific manner. Transgene expression peaks 3-4 weeks after AAV microinjection and is relatively stable in the following weeks. AAVs (DJ serotype) were generated in HEK293T cells with the AAV-DJ Helper Free Packaging System (Cell Biolabs, Inc., cat# VPK-400-DJ) using the manufacturer-suggested protocol. Viral particles were purified using Virabind AAV Purification Kit (Cell Biolabs, Inc., cat# VPK-140). AAV titers were approximately 1.0 ×10^11^ infectious units/ml. The following AAV constructs were used:

*AAV(DJ)-EF1a-DIO-eNpHR 3.0-EYFP (AAV-DIO-eNpHR).* We used AAV-DIO-eNpHR to inhibit CA3 or MEC neurons projecting to CA1. This virus is double floxed and requires Cre to be expressed. Cre was delivered by rgAAV-Cre which allowed for circuit-specific expression of eNpHR 3.0 and red light neuronal inhibition (660 nm).

*AAV(DJ)-EF1a-DIO-EYFP (AAV-DIO-EYFP).* We used AAV-DIO-EYFP as a Cre-dependent control virus with no opsin or DREADD. This virus is double floxed and requires Cre to be expressed. Cre was delivered by rgAAV-Cre which allowed for circuit-specific labelling with eYFP.

*AAV(DJ)-hSyn-DIO-hM4D(Gi)-mCherry (AAV-DIO-hM4D_i_).* We used AAV-DIO-hM4D_i_ to inhibit CA3 or MEC neurons projecting to CA1. This virus is double floxed and requires Cre to be expressed. Cre was delivered by rgAAV-Cre which allowed for circuit-specific expression of hM4D(Gi) and inhibition mediated by the DREADD agonist C21.

*AAV2-retro helper-CaMKII-mCherry-P2A-Cre (rgAAV-Cre).* rgAAV-Cre infects cells retrograde to the site of injection and expresses Cre recombinase. We used rgAAV-Cre in combination with Cre-dependant viruses to achieve precise circuit specific viral expression. rgAAV-Cre was prepared at a 4× concentration to optimize retrograde transport.

*AAV1-hSyn-Cre-WPRE-hGH (AAV1-Cre).* AAV1-Cre was purchased from Addgene (Addgene#105553-AAV1; pENN.AAV.hSyn.Cre.WPRE.hGH). It produces large quantities of Cre recombinase in the synaptic terminals of infected neurons, some of which gets transported anterogradely to the post-synaptic neurons^32^. This virus was injected into CA3 or MEC in Rosa-YFP mice to label postsynaptic target cells in CA1.

*AAV(DJ)-CaMK2α-NpACY (AAV-NpACY).* We used AAV-NpACY to inhibit CA3 neurons projecting to CA1 in PV-Cre mice and to inhibit individual hippocampal subregions in WT mice. AAV-NpACY contains both enhanced channelrhodopsin-2 (ChR2-H134R) fused to enhanced yellow fluorescent protein (eYFP) and halorhodopsin 3.0 (NpHR3.0). These opsins are spectrally compatible, allowing for neuronal excitation by ChR2 with blue light (473 nm) and neuronal silencing by NpHR3.0 with red light (660 nm). In this experiment only red light inhibition was used. Expression of NpACY in principal neurons was driven by the CaMK2α promoter.

*AAV(DJ)-CMV-GFP (AAV-GFP).* We used AAV-GFP as a control virus for AAV-NpACY. pAAV-CMV-GFP was a gift from Dr. Connie Cepko (Addgene plasmid # 67634; http://n2t.net/addgene:67634; RRID:Addgene_67634). Expression of GFP was driven by the CMV promoter.

*AAV(DJ)-hSyn-DIO-hM3D(Gq)-mCherry (AAV-DIO-hM3D_q_).* We used AAV-DIO-hM3D_q_ to excite PV^+^ interneurons in the CA1 of PV-Cre mice. This virus is double floxed and requires Cre to be expressed. Cre recombinase was expressed in PV^+^ interneurons in PV-Cre mice, limiting the expression of the virus to local CA1 PV^+^ interneurons only.

### Other injectables

*Cholera Toxin subunit-b (CTb).* CTb is a retrograde tracer derived from the non-toxic subunit-b of cholera toxin. For retrograde tracing, mice were unilaterally injected in both CA1 and DG with CTb conjugated with either a 488nm or a 647nm fluorophore. The survival time between injection and perfusion was two days.

### Surgery

Surgeries occurred between 6 and 10 weeks old. Mice were pre-treated with atropine sulfate (0.1 mg/kg, i.p.), anesthetized with chloral hydrate (400 mg/kg, i.p.), administered meloxicam (4 mg/kg, s.c.) for analgesia, and placed into stereotaxic frames. The scalp was incised and retracted, and holes were drilled above the target area. Unless otherwise specified, viruses were injected bilaterally via a glass micropipette connected via polyethylene tubing to a 15 microsyringe (Hamilton) at a rate of 0.1 μl/min and remained in place for an additional 5 min to ensure virus diffusion. Following microinjections, the scalp was sutured and Polysporin was applied to the wound. Mice were administered 0.9% saline (0.5-1.0 ml, s.c.) and placed in a clean cage on a heating pad to recover.

For all experiments involving virus microinjections, only mice showing strong bilateral expression (i.e., expression limited to the target brain region and observable in a minimum of 3 brain sections) were included in the final data set. Additionally, for optogenetics and calcium imaging experiments, only mice with correct optic fiber or GRIN lens placements were included in the final data set.

*Viral microinjection and fiber implantation.* For the circuit-based optogenetics or chemogenetic experiments, the rgAAV-Cre vector was bilaterally microinjected (0.8 ml/side, 0.1 ml/min) into CA1 (AP -1.8, ML ±1.5, DV -1.5 mm) and/or DG (AP -1.8, ML ± 1.65, DV -2.15) and AAV-DIO-eNpHR, AAV-DIO-hM4D_i_, or AAV-DIO-EYFP were injected into either CA3 (AP -2.2 mm from bregma, ML ±3.15 mm, DV -2.65 mm) or MEC (AP +0.2 mm from transverse sinus, ML ±3.5 mm from lambda, DV -2.2 mm from surface of skull). Optical fibers were constructed by inserting and securing a 10 mm length of optical fiber (200 mm diameter, 0.37 NA) into a 1.25 mm zirconia ferrule such that the fiber extended 3 mm beyond the ferrule. These fibers were implanted 0.2 mm dorsal to CA3 or MEC injection site and secured to the skull with dental cement and 2-3 jeweler screws. For single region optogenetic experiments AAV-NpACY or AAV-GFP was injected into either CA1, CA3, DG, or MEC and optic fibers were implanted 0.2 mm dorsal to the injection site. For anterograde tracing, AAV1-Cre was injected in CA3 or MEC. For the combined manipulation of CA3-to-CA1 inhibition with CA1 PV^+^ interneuron excitation, AAV-NpACY or AAV-GFP were injected into CA3 and AAV-DIO-hM3D_q_ was injected in CA1. Mice were allowed to recover for at least 3 weeks while viruses were expressed. *GRIN lens implantation.* Lenses for calcium imaging were implanted as previously described^50,51^. A craniotomy was performed above the right CA1 (AP -2.0 mm, ML +1.5 mm), the dura was carefully removed, and cortical tissue was gently aspirated while continuously applying chilled artificial cerebrospinal fluid. Finally, the microendoscope lens (1-mm diameter, 4-mm length, Inscopix, USA) was implanted at a depth of -1.5 mm relative to the skull surface and fixed to the skull using dental cement and 3 jeweler screws. Then, 2-3 weeks later, the mice were again given anaesthesia to attach the baseplate. The Miniscope (Open-source V4 Miniscopes) attached baseplate was lowered until a well-exposed field of view was observed, then baseplate was fixed with dental cement. After baseplate surgery, mice were habituated microendoscope camera before recording.

### Behaviour

*Fear conditioning.* Contextual fear conditioning occurred in test chambers (31 × 24 × 21 cm) with shock-grid floors (Med Associates). During training, mice were placed in the chambers for 2 min, after which two foot shocks (0.5 mA, 2 s duration, 1 min apart) were delivered. Mice were removed from the chambers 1 min after the last shock (4 min total). The next day, mice were placed in a similar but novel chamber (context B) for 5 min. Depending on the experiment, either 5 hrs or 24 hrs later the mice were placed in the original training chamber (context A) for another 5 min. context B was approximately the same size as context A, with white plastic floor and triangular white plastic walls. Mouse behavior was recorded with overhead cameras and FreezeFrame v.3.32 software (Actimetrics). For contextual fear memory tests, the amount of time mice spent freezing was scored during the entire test session automatically using FreezeFrame software or manually by a researcher blinded to the conditions (for optogenetics experiments). Freezing was defined as the cessation of movement, except for breathing^24^.

*Optogenetic stimulation.* Mice used in the optogenetics experiments were habituated to the optic patch cables for 5 min on the day before training. For optogenetic inhibition during training mice were attached to the patch cables and placed in Context A. For the duration of the training period as described above continuous red light (660 nm, 7-10 mW) was delivered. Freezing behavior during these memory tests was scored manually by an experimenter blinded to the experimental conditions.

*In-vivo calcium activity recording.* We used the UCLA miniscope V4 system (http://www.miniscope.org/) to study neuronal dynamics associated with contextual fear memory in freely behaving mice^33^. Mice were habituated to the miniscope for 15 min/d for 3 d before training. Mice were imaged in the homecage for 5 min 24 h, 3 h and immediately before training. Training consisted of placing mice in the chamber and, 5 min later, delivering two footshocks (0.5 mA, 2 s duration, 1 min apart), and another 5 min later mice were removed from the chamber. 24 h after training, mice were placed in context B for a 10 min test during which the amount of time spent freezing was assessed. 48 h after training mice were placed back in context A for 10 min and freezing was assessed.

### Histology

*Perfusion and tissue preparation.* Mice were transcardially perfused with chilled 1× PBS followed by 4% paraformaldehyde (PFA), fixed in PFA overnight at 4 °C, and transferred to 30% sucrose solution for at least 72 h. When appropriate, perfusions were timed to occur 90 min after training. Brains were sectioned coronally or sagittally using a cryostat (Leica CM1850), and a 1⁄4 sampling fraction was used to obtain 4 sets of 50-μm sections. Sections for immunohistochemistry were stored at 4 °C in PBS containing 0.1% NaN^3^ until staining.

*Immunohistochemisty.* Immunofluorescence staining was conducted as previously described (Ramsaran et al., 2023, KO). For experiments involving quantification of the number of cells positive for immunofluorescence all staining was performed at once using the same antibody solutions. Free-floating sections were blocked with PBS containing 4% normal goat serum and 0.5% Triton-X for 1 h at room temperature. Afterwards, sections were incubated with primary antibodies in fresh blocking solution: rabbit anti-c-Fos (1:500, Synaptic Systems, cat#226003), rat anti-c-Fos (Synaptic Systems, cat #226017), chicken anti-GFP (1:1000, Aves, cat# GFP-1010), rabbit anti-RFP (1:1000, Rockland Immunochemicals, cat# 600-401-379), mouse anti-PV (1:1000, Sigma, cat# 1572), rabbit anti-SST (1:250, Synaptic Systems, cat#257003) for 24 h or 48 h (for c-Fos and PV experiments) at 4 °C. Sections were washed three times for 15 min each with PBS containing 0.1% Tween-20 (PBST), then incubated with PBST containing secondary antibodies: goat anti-chicken Alexa Fluor 488 (1:500, Invitrogen, cat# A-11039), goat anti-mouse Alexa Fluor 647 (1:500, Invitrogen, cat# A-21235), goat anti-rabbit Alexa Fluor 568 (1:500, Invitrogen, cat# A-11011), goat anti-rat Alexa Fluor 633 (1:500, Invitrogen, cat# A-21094) for 2 h at room temperature. Sections were washed with PBS, counterstained with DAPI, mounted on gel-coated slides, and coverslipped with Permafluor mounting medium (ThermoFisher Scientific, cat# TA-030-FM).

*Imaging.* Images were obtained using a confocal laser scanning microscope (LSM 710; Zeiss). For visualization of virus expression, images were acquired with a 20× objective. For image quantification, z-stacks were acquired using a 40× objective (N.A. = 1.3; 15-40 slices with a 1 μm step size). For all experiments involving quantification of the number of cells positive for immunofluorescence all images were acquired using identical imaging parameters (laser power, photomultiplier gain, pinhole, and detection filter settings) in a minimal number of imaging sessions (when possible, in one session). For each experiment, imaging parameters were set using a sample section from a control mouse.

*Quantification.* For cell counting experiments in CA1 every fourth section was assessed for the marker(s) of interest. For each mouse, cells were counted manually in Fiji (National Institutes of Health) using 5 images acquired from 3-5 sections and averaged. To estimate the number of DAPI^+^ cells in CA1 pyramidal layer, DAPI^+^ cells were counted in a small volume (approximately 12-20 ×10^3^ 𝜇m^3^) to obtain the DAPI^+^ density (mm^3^). For all experiments, the volume of the pyramidal layer within each image was measured and multiplied by the DAPI^+^ density value for the appropriate age group to obtain the estimated DAPI^+^ cell number. The proportion of non-PV^+^ pyramidal layer cells (putative excitatory population) expressing c-Fos or YFP are reported as P(c-Fos^+^|PV^-^) or P(YFP^+^|PV^-^) and the proportion of PV^+^ cells expressing c-Fos or YFP are reported as P(c-Fos^+^|PV^+^) or P(YFP^+^|PV^+^).

### Calcium imaging

*Pre-processing*. Raw images were acquired at 20 frames per second and downsampled to 10 frames per second by frame averaging. Raw videos were concatenated and motion corrected with NoRMCorre^52^. Calcium traces and spatial footprints were extracted from the video data using CNMF-E using the previously described parameters^51,53^.

*Spontaneous data removal*. Before further analyses, spontaneous neural activity was removed from the calcium traces recorded during training and test sessions in order to reveal task-specific activity. This method, adapted from Stringer et al. (2019)^15^, involved determining each neuron’s spontaneous activity by concatenating its activity traces from three home cage sessions. The mean and standard deviation of this spontaneous activity trace was then used to z-score each neuron’s activity, both during the home cage as well as the experimental sessions. Subsequently, Principal Component Analysis (PCA) was applied to the population’s spontaneous activity to identify the top 50 components of correlated activity. Finally, each session’s activity was projected along these identified directions of spontaneous activity variation, and the resulting components were subtracted to eliminate correlation statistics shared between the home cage and experimental sessions.

*Absolute pairwise correlation*. For each training and testing session, we computed the neuron-by-neuron Pearson correlation matrix using the cleaned calcium traces. We then took the absolute value of each correlation coefficient and averaged across all unique neuron pairs (excluding the diagonal) to obtain a single measure for each mouse.

*Eigenspectrum analysis*. For each session’s activity, the population’s cleaned calcium traces were organized into two matrices, X and X1. X contained all time points except the last, while X1 contained all time points except the first, such that each row of X and X1 represented a state and next-state pair respectively. Assuming a discount factor (gamma) of 0.9, matrix A was calculated as the following:

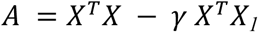

Thereafter, the eigendecomposition of A yielded complex eigenvalue and eigenvector pairs. For each complex eigenvalue, its conjugate was identified, and the real part of their value was recorded. This sequence of real parts of eigenvalues was then sorted and designated as the TD-eigenspectrum for that session’s activity. We observed that this TD-eigenspectrum also followed a power-law, and its power-law decay was measured. Additionally, the effective rank of this set of (real part of) eigenvalues was also computed.

### Statistical Analyses

No statistical tests were used to predetermine sample sizes, but our sample sizes were similar to those reported in previous publications^7,25,51,54^. Mice were pseudo-randomly assigned to all groups to achieve roughly equivalent group sizes. During data collection and quantification, experimenters were blinded to group assignment, except for testing context during data collection and testing context during quantification.

Data were analyzed using paired t-tests, unpaired t-tests, or analysis of variance (ANOVA) with repeated measures when appropriate. When appropriate, ANOVAs were followed by Tukey’s post-hoc comparisons. In some experiments control mice were pooled into a single control group. Sub-group means were not statistically different from one another (data not shown). Both male and female mice were used in all experiments, and no differences were found between the two groups. Statistical significance was set at *P* < 0.05, and Bonferroni correction was applied when appropriate. Statistical analyses were performed Graphpad Prism (version 10.0.1) or using customs scripts in R.

### Computational model

The computational modelling setup followed the experimental setting. An agent’s trajectory in two distinct environments, each with a different valence, was simulated: one (analogous to the shock context) where the agent received a feedback numerical value at the end of the trajectory and the other (analogous to the neutral context) where the agent received no feedback. The feedback numerical value received by the agent is referred to as “reward” in the reinforcement learning literature. Each state visited by the agent, si, was represented by an n-dimensional feature vector. The agent was tasked with learning a linear mapping, parameterized by an n-dimensional vector, w, from each state’s representation to a scalar value indicative of the expected cumulative rewards from that state onward. To achieve this, the agent minimized the temporal difference (TD) error.

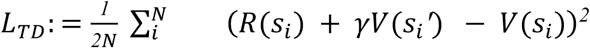

where 𝑠_*i*_, 𝑠_*i*_′ denote the agent’s current and next states respectively, 𝑅(𝑠_*i*_) denotes the reward received by the agent from the environment at state 𝑠_*i*_, 𝛾 is the discount factor that controls the weighting of future rewards for determining the value of the current state, and 𝑉(𝑠_*i*_) denotes the value function estimate of state 𝑠_$_. Note that we define 𝑉(𝑠_*i*_) ∶= 𝑤^*T*^𝑓(𝑠_*i*_) where 𝑓(𝑠_*i*_) is the n-dimensional representation of state 𝑠_*i*_.

Training involved minimizing the TD error by updating 𝑤 from a single trajectory in each environment, with both trajectories containing an equal number of visited states. This setup mirrored an animal spending an equivalent amount of time in each environment. In our model, the number of states in each environment were set to be 20 and n = 40, i.e. each state representation to be a 40-dimensional vector.

Following training, the discrimination ratio was computed to quantify the agent’s ability to differentiate between the rewarding and non-rewarding environments. This was done by using the mean value function estimates across all states in each environment as a proxy for the net value of that environment.

To investigate the relationship between the spectral properties of the state representation space and the agent’s discrimination ability, this setup was run multiple times, each with different properties of the state representation space. Specifically, the eigenspectrum of the representation space was modulated in each run in the following way. All state representations from both environments were concatenated to form the state representation matrix, 𝑋_*all*_. This matrix was then organized into 𝑋 and 𝑋_*1*_, where 𝑋 contained all time points except the last, and 𝑋_*1*_ contained all time points except the first. In this way, each row of 𝑋 and 𝑋_*1*_ represented a state and its corresponding next state, respectively.

Assuming a discount factor (𝛾) of 0.9, matrix 𝐴 was constructed as:

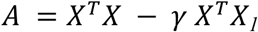

Using gradient descent, the state representations in 𝑋_*all*_ were optimized to minimize the following loss function, resulting in a power-law distribution in the eigenspectrum of matrix 𝐴 with a desired decay coefficient, 𝛼.

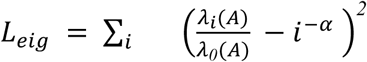

where 𝜆_1_(𝐴) denotes the 𝑖*^th^* eigenvalue of A (arranged in a decreasing order).

For each optimized *X_all_* matrix, the power-law decay coefficient and effective rank metrics were computed using the eigenspectrum of the corresponding 𝐴 matrix. Finally, these spectral metrics were plotted against the agent’s discrimination ratio to characterize the relationship between the dimensionality of the representation space and behavioral performance.

### Analytical Justification

In this section, we provide an analytical justification for using matrix A to characterize the dimensionality of the representation space.

We consider a reinforcement learning problem where an agent interacts with an environment. The agent’s goal is to learn a value function, 𝑉(𝑠), that estimates the expected future reward from a given state, s. The value function is parameterized by a vector, 𝜔, such that for a state 𝑠_*i*_ with corresponding n-dimensional feature vector, 𝑥_*i*_, the value is estimated as the following.

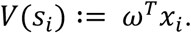

Now, we have a sequence of an observed trajectory that is N-step long, where each step, 𝑖, consists of a state feature vector 𝑥_i_, a next state feature vector 𝑥_i+1_, and a corresponding scalar reward 𝑟_i_ that the agent receives from the environment at step 𝑖. The learning process involves updating the parameter vector 𝜔 to minimize the temporal difference (TD) error.

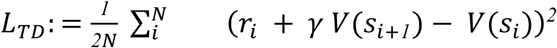

where 𝛾 is the discounting factor that determines how strongly the future rewards are considered while estimating the value function of the current state. Incorporating the linear assumption for the value function,

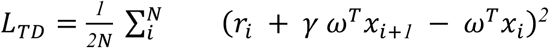

Note that this is a bootstrapped learning problem, and we will use gradient descent to update 𝜔. While computing the gradients, as commonly done in bootstrapped learning problems, the target is assumed to be held fixed and gradients are not computed. To indicate this, we add a stop-gradient operator, □, in the TD-loss formulation.

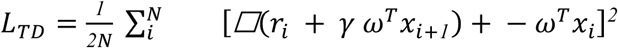

To simplify the expression, we will use a vectorized notation of the TD-loss. Let 𝑋 and 𝑋_*1*_be matrices that contain the concatenated vectors 𝑥_i_ and 𝑥_i+1_, respectively, and 𝑅 be a vector containing the rewards. Note that 𝑋, 𝑋_*1*_ ∈ *ℜ*^Nxn^, and 𝑅, 𝜔 ∈ *ℜ*^n^.

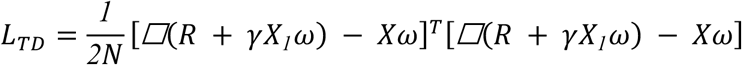

Computing the gradient of the TD-loss with respect to 𝜔:

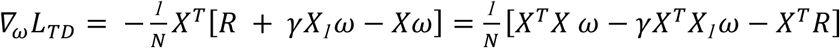

Therefore, the update for 𝜔, written as a continuous time differential equation, is as follows.

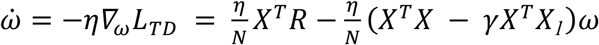

Let us denote 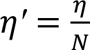 as the effective learning rate, 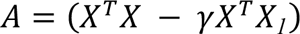 and assume that 𝐴 is diagonalizable as 𝐴 = 𝑃𝐷𝑃^-^*^1^*, where 𝑃, 𝐷 can be complex-valued matrices. Let us also assume that 𝜔 = 𝑃𝑍. Since 𝐴 is constant, i.e. does not change through training, 𝑃 and 𝐷 are also constants.

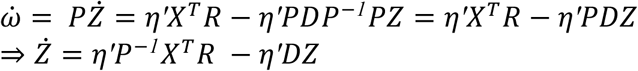

Now, 𝑃^-^*^1^*𝑋^T^𝑅 is a 𝑁 × 𝑛 matrix that does not change over training. Let us denote it as 𝐵.

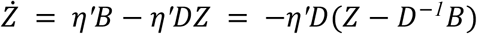

Since 𝐷^-*1*^𝐵 is constant over training, we can denote 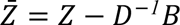 such that 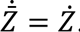.

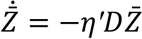

This is a simple first-order linear ordinary differential equation for each component, which has the solution 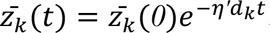. This expression shows that each component of the transformed variable 𝑍 decays exponentially, and the rate of decay is governed by the corresponding eigenvalues of 𝐴, 𝑑_k_. If 𝐴 is low-dimensional and has many dimensions where 𝑑_5_ is small, the dynamics of learning will be slow and thereby it will take many iterations for the value function to converge to a good approximation. In summary, the spectrum of 𝐴 controls the learning dynamics of 𝜔 and subsequently, the agent’s downstream reinforcement learning behavior. For a more detailed description of the relationship between the spectral metrics and (training and generalization) behavior, we defer the reader to Agrawal et al. 2022^55^ and Ghosh et al. 2022^56^.

### Reporting summary

Further information on research design is available in the Nature Research Reporting Summary linked to this article.

## Data availability

Data are available from the corresponding author on request.

## Code availability

Code is available from the corresponding author on request.

## Acknowledgments

We thank members of the J/F laboratories for their discussions, feedback, and support throughout the work and A. DeCristofaro, D. Lin, M. Yamamoto and W. He for technical assistance. This work was funded by Brain Canada (P.W.F., S.A.J.), Canadian Institutes of Health Research (CIHR) grants PJT180530 and PJT180538 (P.W.F.), Natural Sciences and Engineering Research Council of Canada (NSERC) Discovery Grants (RGPIN-2022-03520 to P.W.F. and RGPAS-2020-00031 to B.A.R.), NSERC Discovery Accelerator Supplement RGPAS-2020-00031 (B.A.R.), CIFAR Child and Brain Development program (P.W.F.), CIFAR Learning in Machines and Brains Program (B.A.R.), Canada AI Chair (B.A.R.), Ontario Graduate Scholarship (C.M.), SickKids Restracomp Fellowship (C.M., A.I.R. and B.W.), NIH grant F31 MH120920 (A.I.R.), Canadian Neurodevelopmental Research Training Platform (A.I.R.), Japan Society for the Promotion of Science KAKENHI (S.S.) and a Vanier Canada Graduate Fellowship (A.G.).

## Author contributions

C.M., A.G., A.I.R., B.A.R., and P.W.F. conceived the study, designed the experiments, and wrote the original manuscript. C.M., A.I.R., and T.L. performed the experiments. A.G., C.M., and B.W. performed the analyses. S.S. contributed to the experimental settings. S.A.J. provided critical feedback on the manuscript.

## Competing interests

The authors declare that there are no competing interests.

## Correspondence and requests for materials

should be addressed to Paul W. Frankland or Blake A. Richards.

---

**Extended Data Fig. 1.**
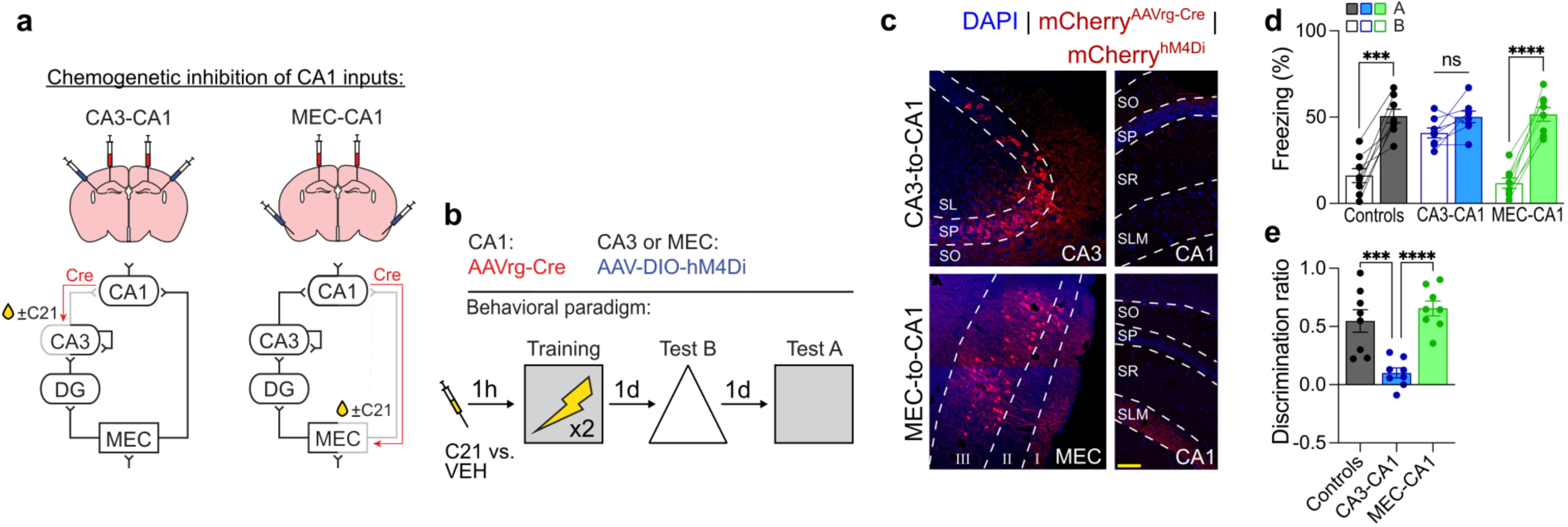
Chemogenetic inhibition of CA3-to-CA1 projections reduces mnemonic resolution. **a)** Experimental design for contextual fear experiments with CA3-to-CA1 or MEC-to-CA1 projection inhibition. AAVrg-Cre was injected into CA1 for all groups and AAV-DIO-hM4D_i_ was injected into either CA3 (trisynaptic expression of eNpHR3.0) or MEC (monosynaptic expression). **b)** C21 or VEH was injected an hour before training and mice were fear conditioned. **c)** Representative images of AAV-DIO-hM4D_i_ expression in CA3 and MEC layer 3 with axons in CA1. **d)** Inhibiting CA3-to-CA1 projections resulted in equivalent freezing in contexts A and B (two-way repeated measures ANOVA; Group x Context interaction: *F*_2,21_ = 14.18, *P* < 0.0001; n = 8 mice per group). **e)** CA3-to-CA1 inhibition reduced memory resolution (one-way ANOVA; Group: *F*_2,21_ = 16.82, *P* < 0.0001; n = 8 mice per group). Scale bars: yellow, 100 μm. Data are shown as mean ± SEM. \**P* < 0.05, \*\**P* < 0.01, \*\*\**P* < 0.001, \*\*\*\**P* < 0.0001. Lines link freezing scores across tests in the same mice.

**Extended Data Fig. 2.**
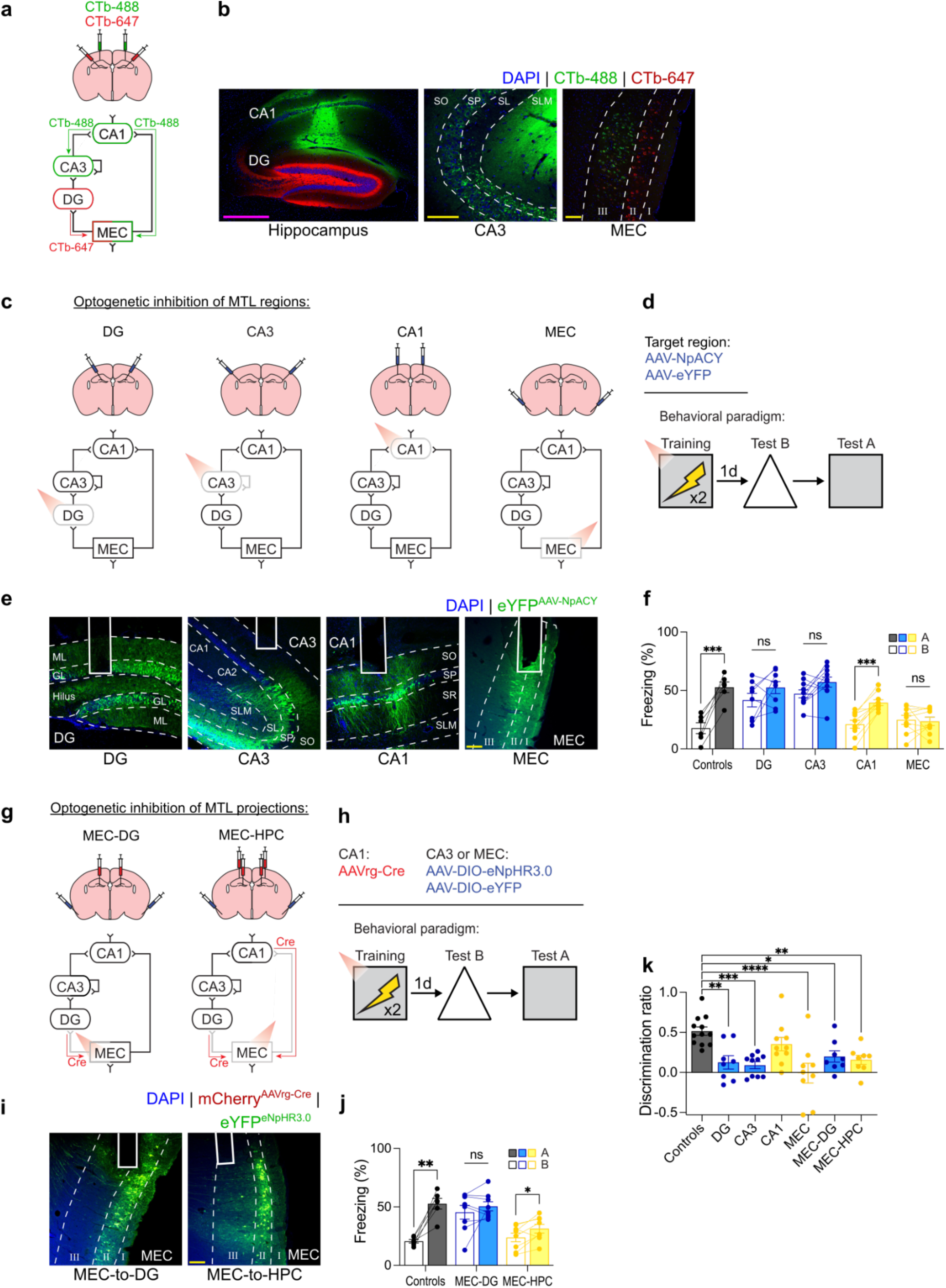
The trisynaptic pathway is necessary for high resolution memories. **a)** Experimental design for retrograde tracing with cholera toxin subunit B. Mice were injected with CTb containing two different fluorophores (CTb-488 and CTb-647) in CA1 and DG and expression was evaluated 2 days post-injection. **b)** Representative images showing CTb injections in CA1 and DG with retrogradely-labeled neurons in CA3 and MEC layer 3 (after CA1 injections) and in MEC layer 2 (after DG injections). **c)** Experimental design for optogenetic inhibition of different regions. AAV-NpACY or AAV-eYFP was injected into either DG, CA3, CA1, or MEC and an optic fiber was implanted above the same region. **d)** Mice were contextually fear conditioned with red light inhibition during the training session. **e)** Representative images of AAV-NpACY expression with optic fiber placement. **f)** Inhibition of DG, CA3, or MEC impaired memory resolution (two-way repeated measures ANOVA; Group x Context interaction: *F*_4,38_ = 6.59, *P* = 0.0004; n = 6 to 10 mice per group). **g)** Experimental design for inhibition of MEC projections. AAVrg-Cre was injected into DG or CA1 and DG for all groups and AAV-DIO-eNpHR3.0 was injected into MEC. Control mice were injected with AAV-DIO-eYFP in MEC. **h)** Contextual fear conditioning with red light inhibition during training. **i)** Representative images showing eNpHR3.0 expression in MEC layer 2 and MEC layers 2 and 3 with optic fibers implanted above. **j)** Inhibition of MEC-to-DG projections reduced memory resolution (two-way repeated measures ANOVA; Group x Context interaction: *F*_2,19_ = 18.29, *P* < 0.0001; n = 6 to 8 mice per group). **k)** Discrimination ratios for region and pathway inhibition experiments (one-way ANOVA; Group: *F*_6,58_ = 6.13, *P* < 0.0001; n = 6 to 10 mice per group). Scale bars: yellow, 100 μm; magenta, 500 μm. Data are shown as mean ± SEM. \**P* < 0.05, \*\**P* < 0.01, \*\*\**P* < 0.001, \*\*\*\**P* < 0.0001. Lines link freezing scores across tests in the same mice.

**Extended Data Fig. 3.**
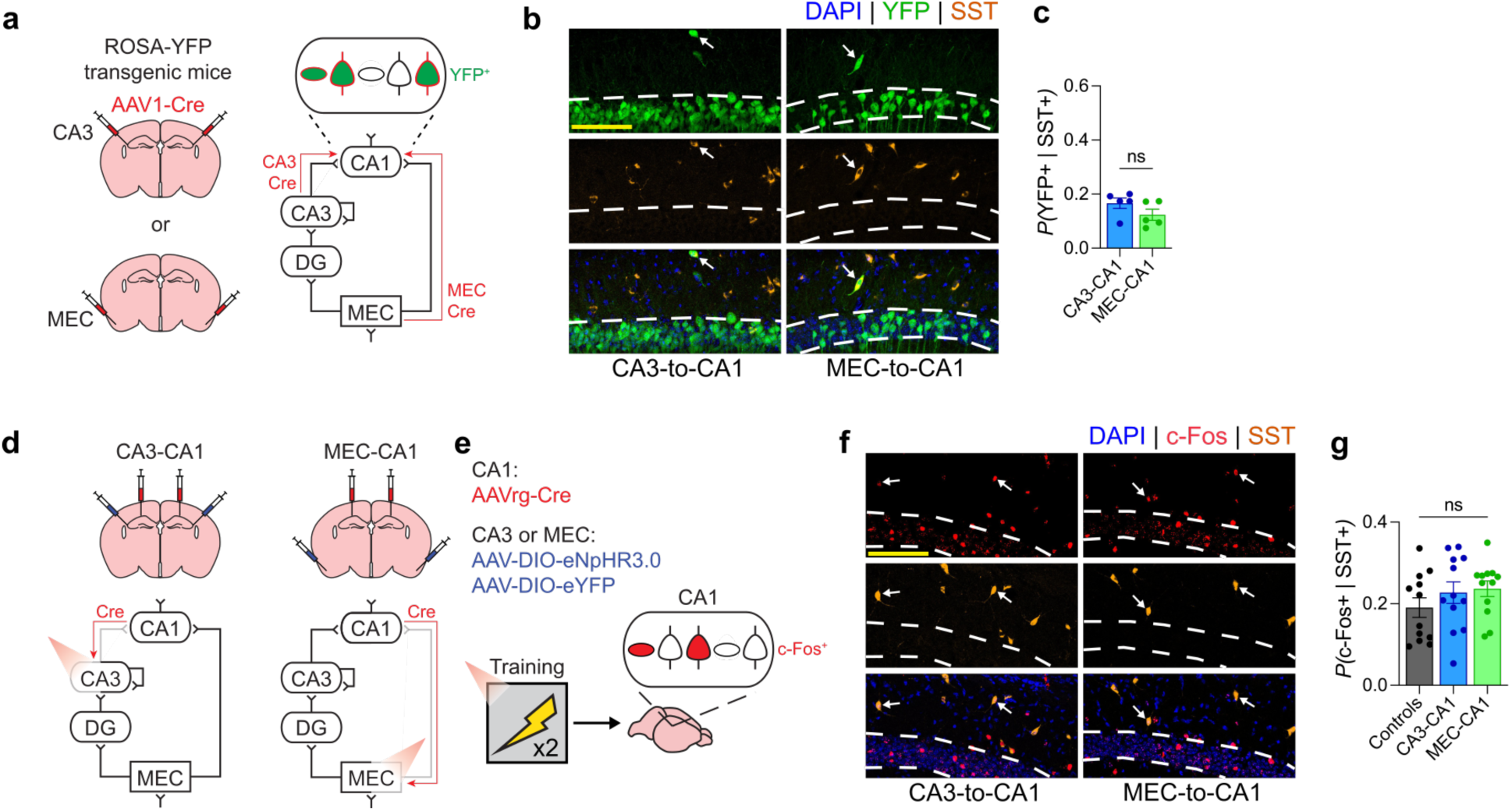
SST^+^ interneurons in CA1 do not receive preferential inputs from either CA3 or MEC. **a)** Transsynaptic anterograde tracing was achieved by injecting AAV1-Cre in CA3 or MEC of ROSA-YFP transgenic mice. Cre protein is transported from the presynaptic to postsynaptic neuron which expresses YFP in the presence of Cre. **b)** Representative images showing YFP and SST co-localization in CA1. White arrows indicate the overlap of YFP and SST. **c)** SST^+^ interneurons in CA1 received similar input from CA3 and MEC (unpaired *t* test; *t*_8_ = 1.49, *P* = 1.1746; n = 5 mice per group). **d** and **e)** Experimental design for examining c-Fos induction following contextual fear training with CA1 input inhibition. Viral strategy and training were the same as previously described (Fig. 2g), mice were perfused 90 min after training and c-Fos in CA1 was quantified. **f)** Representative images of c-Fos and SST co-labelling in CA1 pyramidal layer. white arrows indicate overlap of c-Fos and SST. **g)** All groups had comparable overlap of c-Fos in SST^+^ interneurons (one-way ANOVA; Group: *F*_2,33_ = 1.09, *P* = 0.3492; n = 12 mice per group). Scale bars: yellow, 100 μm. Data are shown as mean ± SEM. \**P* < 0.05, \*\**P* < 0.01, \*\*\**P* < 0.001, \*\*\*\**P* < 0.0001.

**Extended Data Fig. 4.**
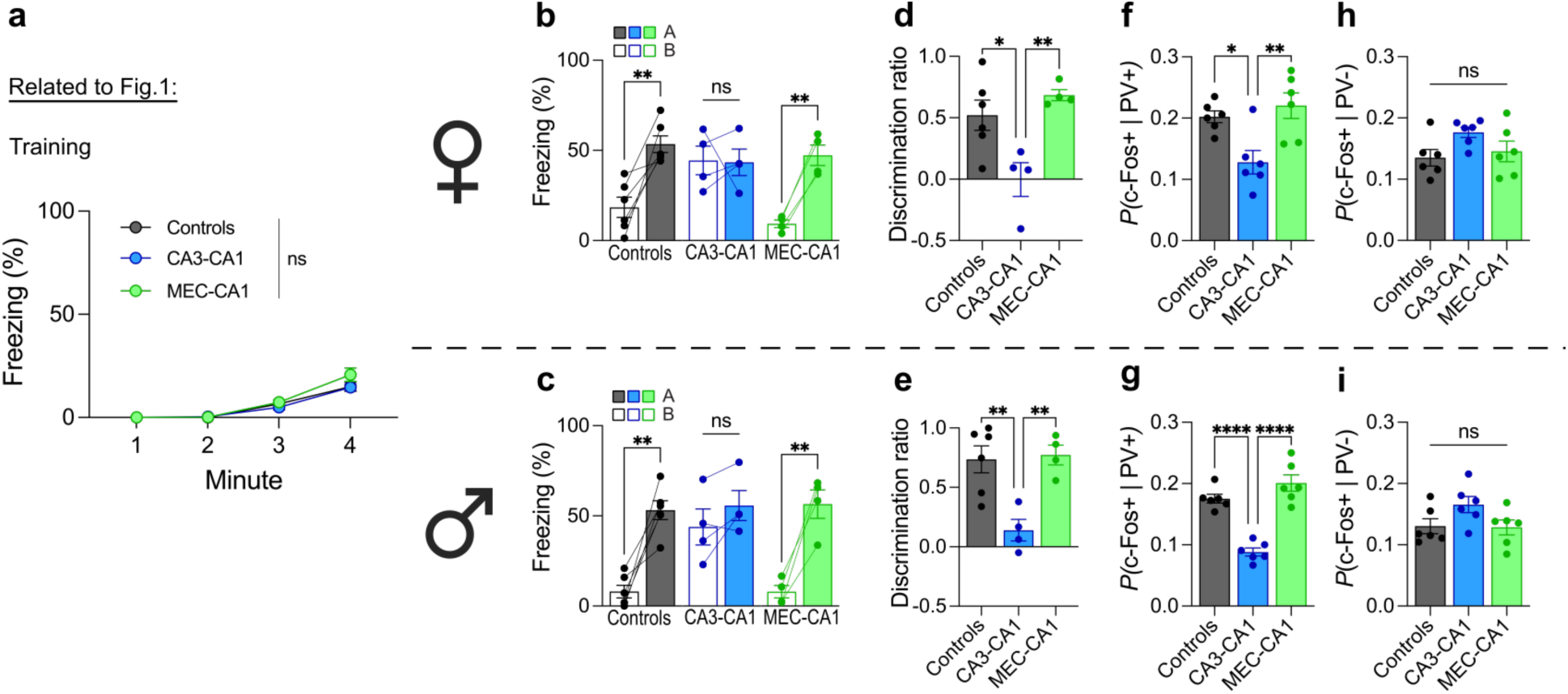
Inhibition of projections to CA1 has no effect on memory acquisition and shows no sex differences. **a)** Inhibiting CA3 or MEC projections to CA1 did not affect fear learning (two-way repeated measures ANOVA; Group x Time interaction: *F*_6,75_ = 1.42, *P* = 0.2194; n = 8 or 12 mice per group). **b)** Inhibition of CA3-to-CA1 projections resulted in equivalent freezing in contexts A and B for female mice (two-way repeated measures ANOVA; Group x Context interaction: *F*_2,11_ = 6.36, *P* = 0.0146; n = 4 or 6 mice per group). **c)** Inhibition of CA3-to-CA1 projections resulted in equivalent freezing in contexts A and B in male mice (two-way repeated measures ANOVA; Group x Context interaction: *F*_2,11_ = 6.50, *P* = 0.0137; n = 4 or 6 mice per group). **d)** Inhibition of CA3-to-CA1 projections reduced mnemonic resolution in female mice (one-way ANOVA; Group: *F*_2,11_ = 8.24, *P* = 0.0065; n = 4 or 6 mice per group). **e)** Inhibition of CA3-to-CA1 projections reduced mnemonic resolution in male mice (one-way ANOVA; Group: *F*_2,11_ = 10.29, *P* = 0.0030; n = 4 or 6 mice per group). **f)** Inhibition of CA3-to-CA1 projections reduced training-induced c-Fos in PV^+^ interneurons in CA1 in female mice (one-way ANOVA; Group: *F*_2,15_ = 7.96, *P* = 0.0044; n = 6 mice per group). **g)** Inhibition of CA3-to-CA1 projections reduced c-Fos in PV^+^ interneurons in CA1 in male mice (one-way ANOVA; Group: *F*_2,15_ = 38.28, *P* < 0.0001; n = 6 mice per group). **h)** Inhibiting projections to CA1 did not effect c-Fos in PV^-^ neurons in CA1 in female mice (one-way ANOVA; Group: *F*_2,15_ = 2.63, *P* = 0.1052; n = 6 mice per group). **i)** Inhibiting projections to CA1 did not effect c-Fos in PV^-^ neurons in CA1 in male mice (one-way ANOVA; Group: *F*_2,15_ = 2.78, *P* = 0.0929; n = 6 mice per group). Data are shown as mean ± SEM. \**P* < 0.05, \*\**P* < 0.01, \*\*\**P* < 0.001, \*\*\*\**P* < 0.0001. Lines link freezing scores across tests in the same mice.

**Extended Data Fig. 5.**
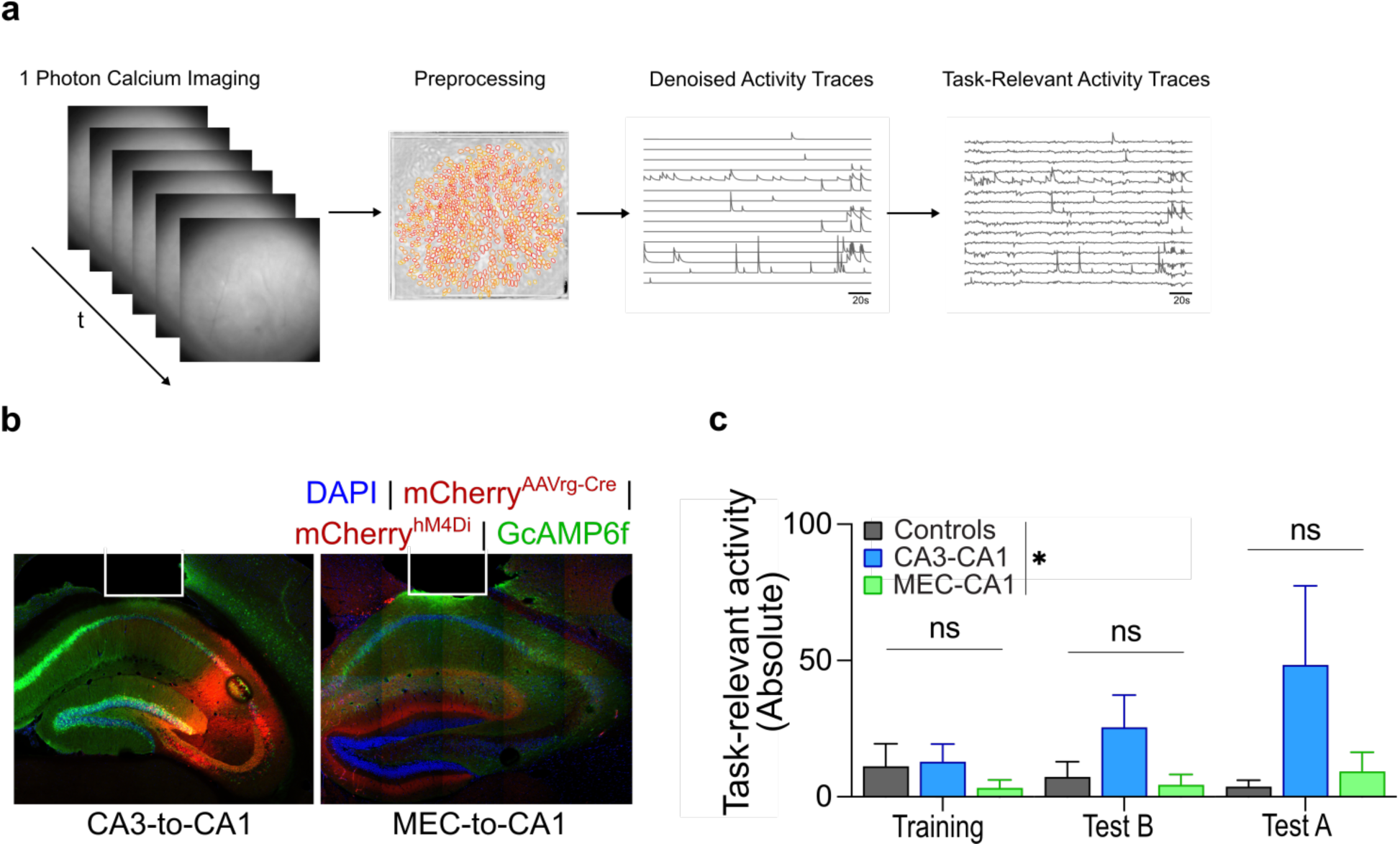
Imaging of CA1 calcium activity with input inhibition during contextual fear conditioning. **a)** Schematic showing the pipeline of recording and processing of 1 photon calcium imaging. **b)** Representative histology showing lens implantation above CA1, GcAMP6f expression (green), and AAV-DIO-hM4D_i_ (red) expression in CA3 (left) or in axons from MEC (right). **c)** Chemogenetically inhibiting projections to CA1 results in a main effect of inhibition on task-relevant activity, however there is no interaction between inhibition and behaviour session (two-way repeated measures ANOVA; Group *F*_2,19_ = 3.77, *P* = 0.0419; Group x Context interaction: *F*_4,38_ = 0.78, *P* = 0.5446; n = 7 or 8 mice per group). Data are shown as mean ± SEM. \**P* < 0.05, \*\**P* < 0.01, \*\*\**P* < 0.001, \*\*\*\**P* < 0.0001.

**Extended Data Fig. 6.**
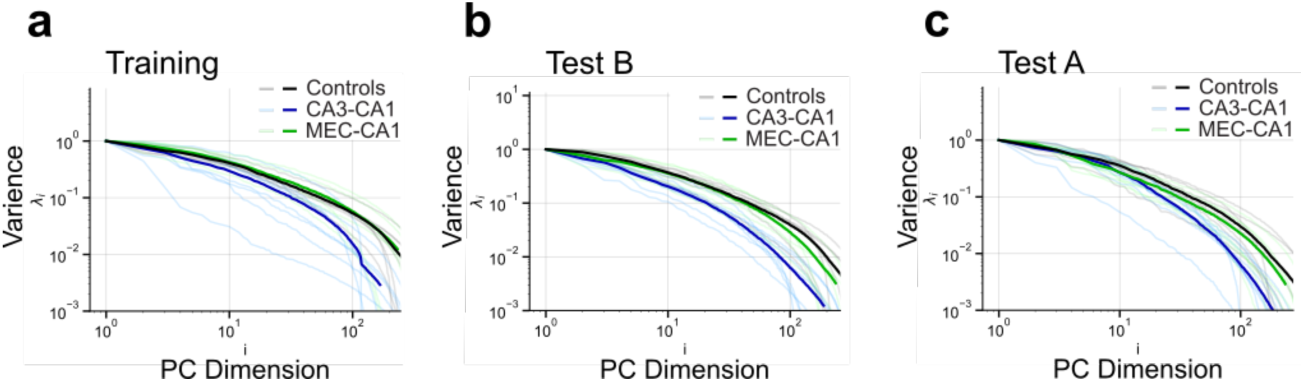
CA1 Eigenspectrum decay is more pronounced with CA3-to-CA1 projection inhibition during contextual fear conditioning. **a)** Eigenspectrum plot for individual mice (faded lines) and group averages (solid lines) in the training session. **b)** Eigenspectrum plot for individual mice (faded lines) and group averages (solid lines) in context B. **c)** Eigenspectrum plot for individual mice (faded lines) and group averages (solid lines) in context A.

**Extended Data Fig. 7.**
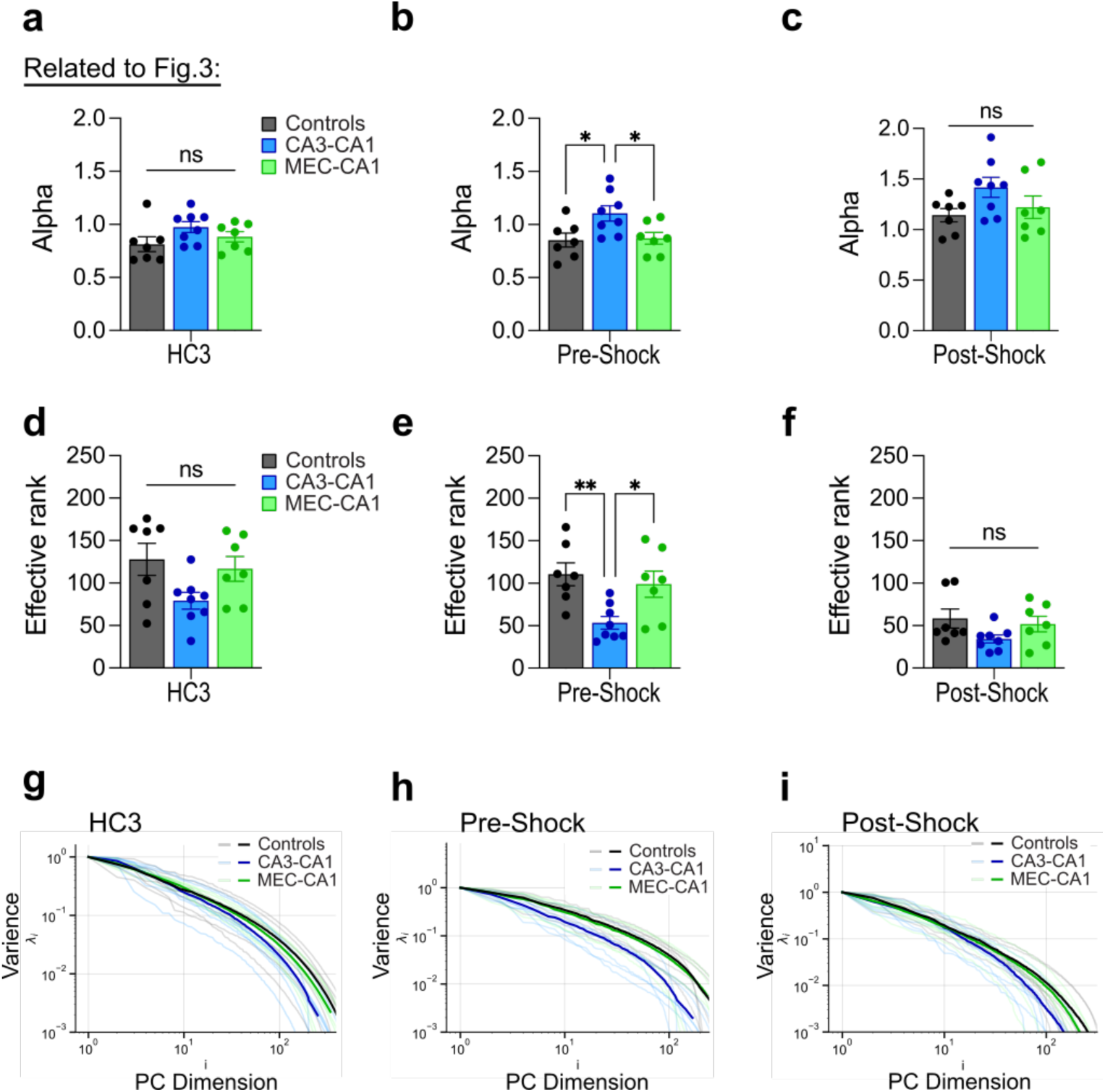
The pre-shock period, but not homecage session 3 or the post-shock period, shows reduced dimensionality following CA3-to-CA1 projection inhibition. **a)** There is no group difference in alpha values during the homecage 3 (HC3) session (one-way ANOVA; Group: *F*_2,19_ = 2.04, *P* = 0.1573; n = 7 or 8 mice per group). **b)** Chemogenetic inhibition of CA3-to-CA1 projections results in higher alpha values for the pre-shock period of the training session (one-way ANOVA; Group: *F*_2,19_ = 4.79, *P* = 0.0206; n = 7 or 8 mice per group). **c)** There is no group difference in alpha values during the post-shock period of the training session (one-way ANOVA; Group: *F*_2,19_ = 2.31, *P* = 0.1266; n = 7 or 8 mice per group). **d)** There is no group difference in rank during the HC3 session (one-way ANOVA; Group: *F*_2,19_ = 3.21, *P* = 0.0628; n = 7 or 8 mice per group). **e)** Chemogenetic inhibition of CA3-to-CA1 projections results in lower rank for the pre-shock period of the training session (one-way ANOVA; Group: *F*_2,19_ = 6.36, *P* = 0.0077; n = 7 or 8 mice per group). **f)** There is no group difference in rank during the post-shock period of the training session (one-way ANOVA; Group: *F*_2,19_ = 2.21, *P* = 0.1370; n = 7 or 8 mice per group). **g)** Eigenspectrum plot for individual mice (faded lines) and group averages (solid lines) in the HC3 session. **h)** Eigenspectrum plot for individual mice (faded lines) and group averages (solid lines) in the pre-shock period. **i)** Eigenspectrum plot for individual mice (faded lines) and group averages (solid lines) in the post-shock period. Data are shown as mean ± SEM. \**P* < 0.05, \*\**P* < 0.01, \*\*\**P* < 0.001, \*\*\*\**P* < 0.0001.

## Notes

### Competing Interest Statement

The authors have declared no competing interest.

